# Functional populations in prefrontal cortex related to working memory encoding and maintenance

**DOI:** 10.1101/2025.03.27.645288

**Authors:** Nicolás Pollán Hauer, Bijan Pesaran, Klaus Wimmer

## Abstract

Nonlinear mixed selectivity, with neurons responding to diverse combinations of task-relevant variables, has been proposed as a key mechanism to enable flexible behavior and cognition. However, it is debated whether the structure of neural population responses in fronto-parietal cortices is better described as random mixed-selective or as non-random, that is, in terms of multiple subpopulations with stereotypical response profiles. Here, we show that neural activity in the macaque prefrontal cortex during a working memory and a visual response task is organized into subpopulations that provide a comprehensive description of the low-dimensional population dynamics. First, analysis of the demixed Principal Components shows that the neural code faithfully represents stimulus identity, task condition, and elapsed time during the trial. Second, a model-free analysis of the population structure reveals a significant degree of clustering, implying a non-random distribution of feature selectivity that is incompatible with random mixed selectivity. Closer inspection shows that stimulus-selective neurons also tend to be task-selective. Third, examining the contribution of stimulus-selective neurons to task-condition-related variance reveals two contrasting activity profiles that correspond to functionally different populations. One population responds during visual stimulation while the other activates during memory maintenance. Finally, the observed neural geometry explains how stable task and stimulus information can be read out from the population response using a linear decoder. Our results highlight that despite the heterogeneity of prefrontal responses during working memory, neurons do not represent random mixtures of task features but are structured according to neural subpopulations.

## Introduction

How does neural activity relate to behavior in cognitive tasks? While much of our understanding of the organization of the early visual system relied on viewing single neurons as feature detectors (***Hubel and Wiesel, 1959***), neurons in the fronto-parietal cortex are best described as mixed selective (***Eichenbaum, 2018***; ***Rigotti et al., 2013***; ***Mante et al., 2013***). The firing rates of mixed selective neurons are modulated by multiple task variables (***Raposo et al., 2014***; ***Bernardi et al., 2020***). This leads to the key question of how information is represented across neurons when considering a population of neurons collectively rather than single neurons in isolation. It is the structure of the neural population activity that ultimately determines the computations underlying behavior.

The structure of the population activity can be classified along two axes, linear vs. nonlinear mixed selectivity and categorical vs. category-free representations (***Kaufman et al., 2022***; ***Tye et al., 2024***). Linear mixed mixed selectivity gives rise to neural representations that are low-dimensional because the coding direction for each variable is independent of coding of the other variables. A consequence of this disentangled representation is that a linear decoder can generalize between task conditions ***Pagan et al. (2013***); ***Bernardi et al. (2020***); ***Nogueira et al. (2023***). Nonlinear mixed selectivity (***Rigotti et al., 2013***; ***Wallis et al., 2001***; ***Mansouri et al., 2006***; ***Dang et al., 2021***), on the other hand, can lead to high-dimensional representations. Those allow for flexibility in decoding, but at the expense of generalization (***Rigotti et al., 2013***). High-dimensional representations arise from category-free mixed selectivity (i.e., random mixed selectivity with no discrete groups of neurons representing particular combinations of task variables). In contrast, in clustered or categorical representations, neurons with similar feature weights cluster together, forming functional subpopulations. The distinction between random mixed selectivity and clustered representations is important because a clustered representation suggests that neurons have special functional roles in the circuit. Taken together, when neural dynamics are low-dimensional, we are justified in searching for functional cell types and thinking about computations in terms of single cells. Previous work has found evidence both for random mixed selectivity (***Raposo et al., 2014***; ***Blanchard et al., 2018***; ***Bernardi et al., 2020***) and for clustered representations (***Hirokawa et al., 2019***; ***Hocker et al., 2021***; ***Yang et al., 2022***).

Here, we investigate the encoding of task variables in the prefrontal cortex during working memory. We test whether the structure of neural population responses in PFC is better described as random mixed-selective or as non-random, that is, in terms of multiple subpopulations with stereotypical response profiles. Our data shows non-random mixed selectivity, as has recently been observed in rodents during simpler working memory tasks (***Hirokawa et al., 2019***; ***Yang et al., 2022***). This suggests a common organizational principle of prefrontal neural activity underlying working memory in primates and rodents.

## Results

### Task structure and encoding of task variables

We analyzed prefrontal single neuron recordings from two adult monkeys that were performing a classical visual working memory task, the memory-guided saccade task and a visual variation of this task (***Figure 1a***; ***Markowitz et al., 2015***). In the memory task, the animals fixate on the screen where a stimulus is presented during 300 ms. The stimuli are presented in one out of eight evenly spaced positions around the fixation point. After the stimulus is switched off, a delay period follows, whose length varies slightly from trial to trial (between 1 and 1.5 seconds). Once the fixation cross is switched off, indicating the end of the delay, the animals report the remembered location with their gaze. In the visual variation of this task (***Figure 1a***, bottom), the cue stimulus is not removed after 300 ms but instead it is maintained until the animals are cued to respond (fixation cross is switched off). The structure of the visual task is very similar to the memory task but it does not require memory. In each recording session, visual and memory trials are randomly interleaved. This task design allows to study single neuron dynamics related to the encoding and maintenance fo working memory (***Markowitz et al., 2015***).

**Figure 1.**
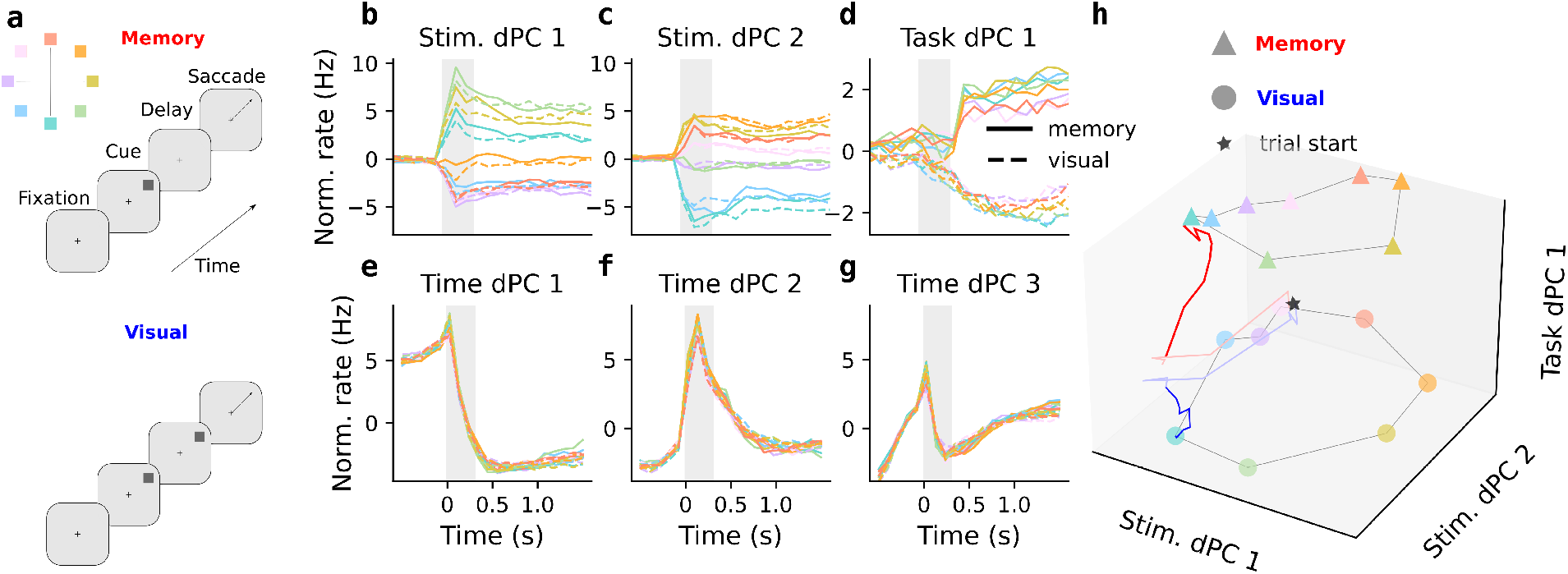
Prefrontal neurons encode time, stimulus location and task condition during a working memory task with a visual variation. **a** Experimental task design. Classic oculomotor delayed response task (top) and a visual variation that does not involve a retention or mnemonic period (bottom). **b-g** Neural trajectories along different demixed-Principal Components illustrate how the different task-relevant variables are encoded in the neural activity. The data were projected onto the respective dPCA decoder axis, so that there are 16 lines 8 cue locations and 2 task conditions. **b**,**c** Neural activity projected along the two first stimulus-related dPCs, each corresponding to one of the orthogonal spatial dimensions of the stimulus. **d** The trajectories along the task condition-related dPC bifurcate when the memory and visual conditions become distinguishable (at the time point corresponding to the stimulus offset in the memory task). **e-g** Activity along time-related dPCs reflects elapsed time and relevant task epochs. **h** The geometry of the neural representation is illustrated by the neural activity in a space spanned by the 2 stimulus dPCs and the task dPC. The star in the middle represents the departing point for all trajectories (any combination of task condition and stimulus location). The colored triangles (circles) correspond to the activity at the end of a memory (visual) for each stimulus location.

We used demixed-PCA (***Kobak et al., 2016***) to understand how different task variables (cue stimulus location, task condition, and elapsed time) are represented in the PFC population activity (***Figure 1b-g***). Together, the 6 dPCs from ***Figure 1*** capture 46.5 % of the total variance, and they allow us to gain an intuition on the neural representation.

As has been observed previously, two stable stimulus components reflect the two-dimensional task geometry, yielding a stable representation of the *x* and *y* coordinates of the visual cue locations (***Figure 1b,c***). This becomes more apparent when plotting the two stimulus components against each other (***Figure 1h***). The neural activity along the task dPC shows that the neural representation during the memory and visual conditions diverges at cue offset (*t* = 300 ms) when the two experimental conditions become distinguishable (***Figure 1d***). The activity along the task dPC (***Figure 1d***) suggests a difference in the neural representation depending on whether memory is needed or not (memory vs visual task). The information in the stimulus and task dPCs can be combined into a three-dimensional representation that reveals the geometry of neural activity (***Figure 1h***). After the 300 ms cue period, the neural activity in both memory and visual conditions follow a ring-like topology of the cue locations (***Figure 1a***). The task-condition related dPC separates both cue representations but when projecting the activity of the memory and visual tasks in a single plane spanned by the stimulus dPCs (horizontal plane in ***Figure 1h***), the responses in the two tasks become indistinguishable. ***Figure 1h*** also illustrates the neural trajectory for one of cues, which first moves in the stimulus direction during the 300 ms cue period before it separates in the task direction. Computationally, this neural geometry suggests that cue location can be decoded independently of the task condition, even though the overall neural code is task-dependent.

Apart from the information related to stimulus location and task condition, a significant proportion of the variance (26.7 %) is captured by dPCs that capture firing rate variations across time that are not stimulus or task-condition related. The three leading time dPCs reflect firing rate changes during all tasks periods, fixation, cue and delay (***Figure 1e-g***). The gradual build-up of activity (ramping activity) before the cue and during the delay (***Figure 1g***) may be related to the tracking of elapsed time (***Gouvêa et al., 2015***), response preparation (***Markowitz et al., 2015***; ***Inagaki et al., 2019***) or urgency signals (***Thura, 2020***; ***Cisek, 2019***).

### Single neuron activity and the structure of selectivity

The activity of the recorded prefrontal neurons spans a spectrum of heterogeneous profiles (***Figure 2a-d***). Some of these firing rate profiles, such as persistent and ramping activity, can be functionally interpreted within the context of the task. For example, the neuron in ***Figure 2a*** shows persistent activity during the cue and delay periods, which has been linked to the maintenance of information and considered as the main neural correlate of working memory (***Fuster and Alexander, 1971***; ***Funahashi et al., 1989***; ***Constantinidis et al., 2001b***; ***Leavitt et al., 2017***; ***Wimmer et al., 2014***). In terms of selectivity, this neuron is described by a large stimulus weight and no selectivity for the task condition, i.e., the activity is highly similar across the two tasks (***Figure 2e***). Another type of firing profile that is amenable to interpretation is a seemingly stimulus-dependent activation, as exhibited by the cell in ***Figure 2b,f***, that resembles the response of neurons in sensory cortices (***Steinfeld et al., 2024***). In addition to a significant stimulus weight, this neuron has a large task weight that accounts for the differential task-dependent activation. The third example neuron, ***Figure 2c,g***, shows ramping activity during delay epochs, but only in the memory task. This has been frequently observed during tasks involving predictable delay lengths (***Inagaki et al., 2019***). Other activity profiles are not easily interpretable, such as the one in ***Figure 2d,h*** that shows transient selectivity in the early delay period, only in the memory task.

**Figure 2.**
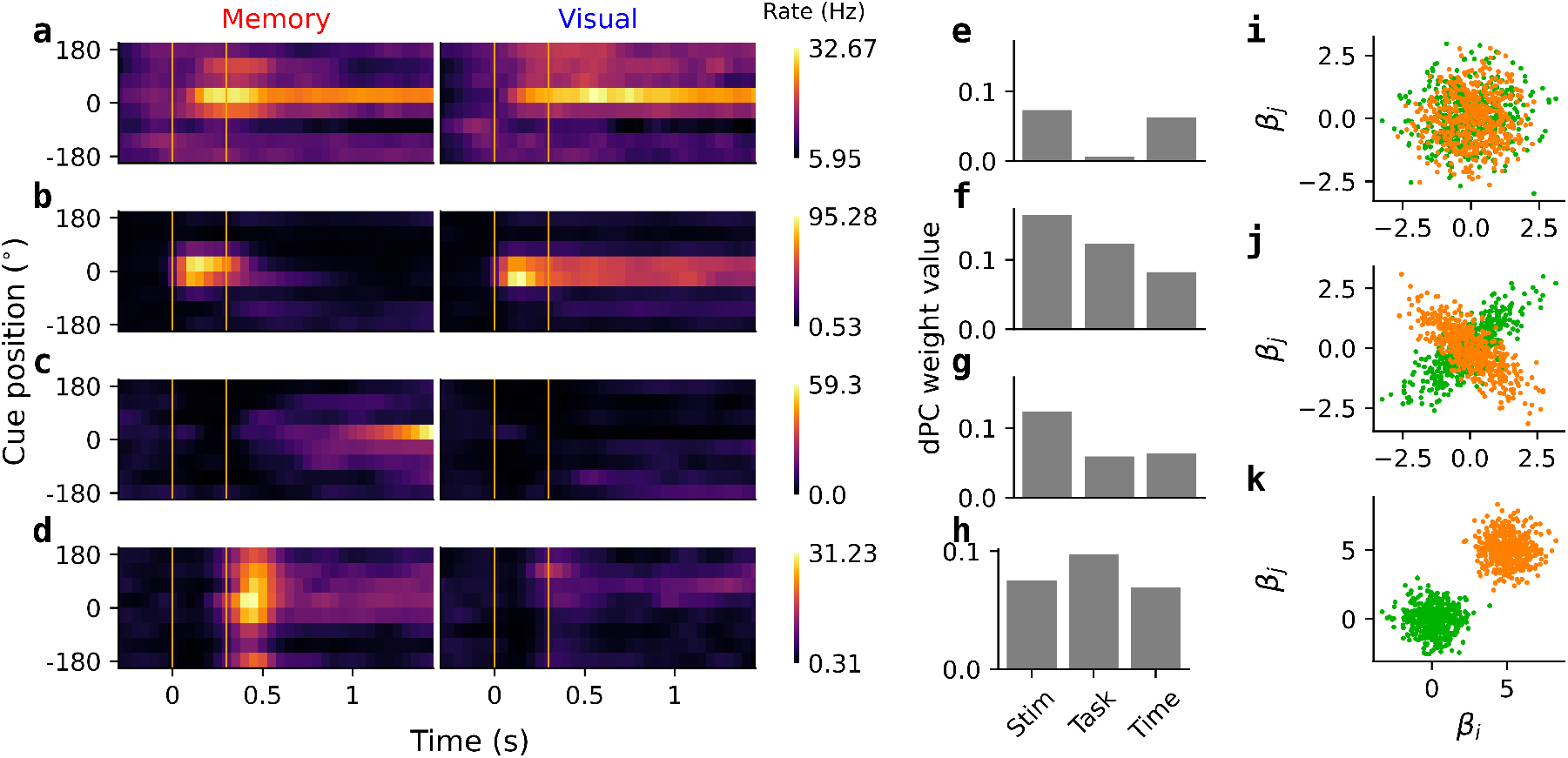
Single neuron activity and the structure of selectivity. **a-d** Average firing rates of four example neurons in response to the eight different cues in the memory and the visual task. **e-h** Contribution of the neurons in **a-d** to different task variables (stimulus location, task condition, and time). **i-k** Cartoon examples illustrating qualitatively different distributions of selectivity among neurons. In **i** the selectivity for variables *β*_*i*_ and *β*_*j*_ is randomly distributed, the orange and green populations are statistically indistinguishable. In **j** the orange and green dots represent neurons with non-random, statistically dependent selectivity. In **k**, neurons form separate clusters, that is, they belong to different functional categories.

Are the neural activity profiles that we observed examples from randomly distributed features among neurons (category free, random mixed selectivity) or are they instead representing different functionally specialized groups of cells? To get an understanding of how the selectivity for the different variables is distributed among the PFC neurons, it is useful to look for structure in a *selectivity-space* (or *feature-space*, where each data point represents a neuron according to its contribution of the different variables considered (cartoon examples in ***Figure 2i-k***). A predominantly random structure (***Figure 2i***) would imply that our example neurons are taken from a continuum of responses, with no underlying structure. A significant degree of structure (***Figure 2j***) would indicate dependencies between the variables and some functional specialization of the neurons. Neurons may also form separate clusters, that is, they could belong to clearly separated functional classes, such as illustrated in ***Figure 2k***. To fully examine how selectivity is distributed across neurons, we moved beyond analyzing only pairs of selectivity weights, and instead considered a higher-dimensional selectivity space and statistical measures at the population level.

### Non-random mixed selectivity

We tested for structure in the distribution of task-related selectivity in the population of neurons in the selectivity space, spanned by task-informative dimensions. In this space, each neuron is represented by a point whose coordinates are given by the weights on the respective dPCs (***Figure 3a***). To test whether neural selectivity shows some degree of clustering or non-random structure, we used a non-parametric test, the test of elliptical projection angle index of response similarity (ePAIRS) (***Hirokawa et al., 2019***; ***Raposo et al., 2014***). This test is based on computing the angular distances between nearest neighbor neurons in the dimensionality-reduced space (spanned by the dPCs in our case) and comparing this original distribution to a null distribution generated by bootstrapping from the original data (***Figure 3b***). The distribution was highly nonrandom (*p* < 10^−9^, ***Figure 3b***), indicating that groups of PFC neurons exhibit similar response profiles.

**Figure 3.**
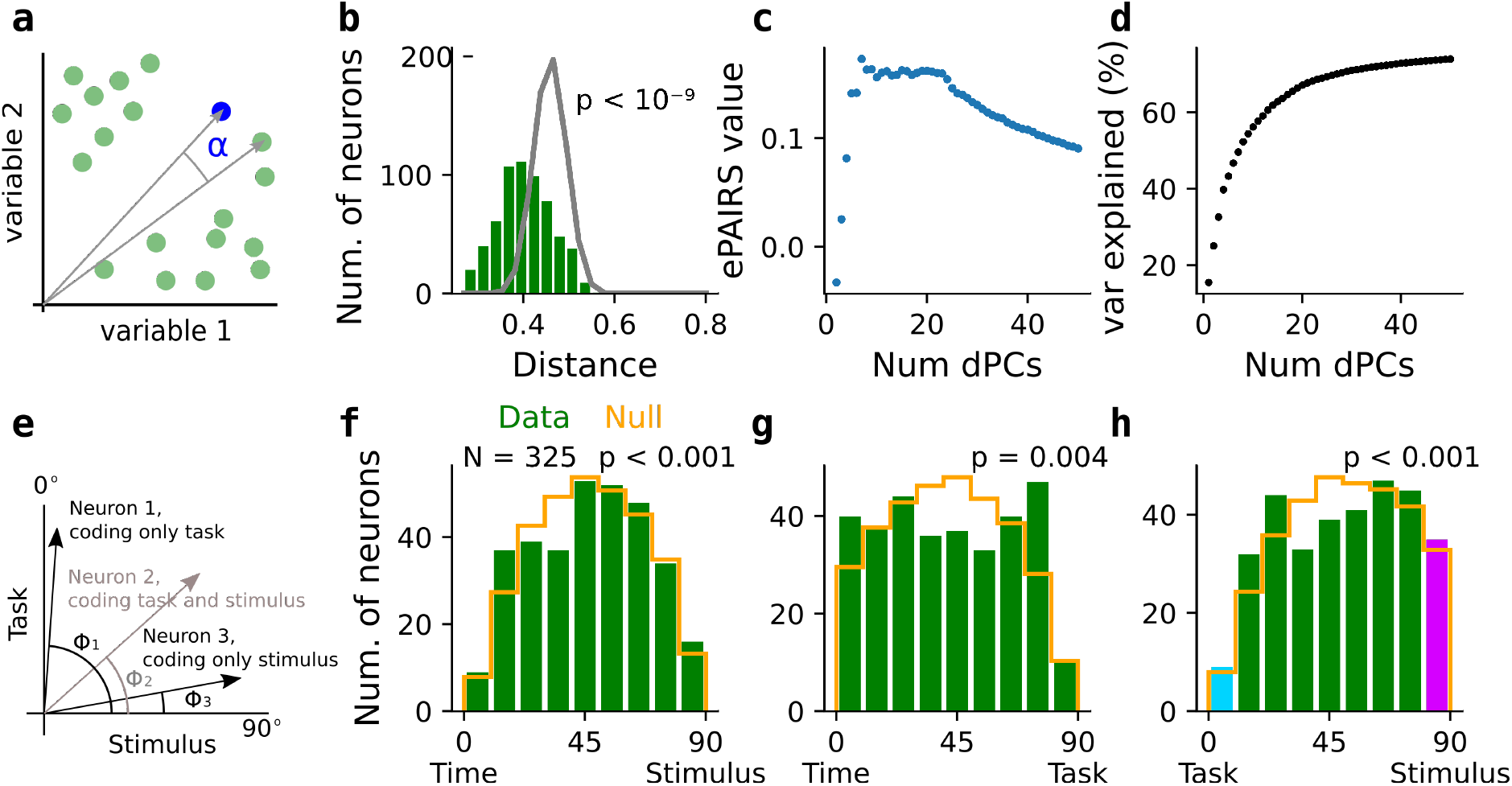
Selectivity for task condition, stimulus location and time is non-randomly distributed among prefrontal neurons. **a-d** Applying elliptical Projection Index of Response Similarity (ePAIRS) to quantified the degree of structure in the distribution of selectivity among prefrontal neurons. **a** Illustration of the ePAIRS test, which is based on calculating the distribution of angular distances between nearest neighbor neurons (green dots) in a space whose axes correspond to the neurons’ selectivity to different task variables. The distribution of angular distances is then compared to a null distribution, obtained by sampling from a multivariate Gaussian distribution with the same covariance as the original data. **b** Distribution of nearest neighbor distances for the data (green histogram) compared to the generated null distribution (gray line) when considering the first 20 dPCs. We used the cosine distance that varies between 0 and 1 as the angular distance varies between 0^°^ to 90^°^. **c** Scalar measure obtained from the ePAIRS test as a function of the number of dPCs that are considered. **d** Variance explained as a function of the number of considered dPCs. **e-h** Distribution of selectivity between pairs of task-variables. **e** Illustration of the analysis: a neuron’s selectivity for a pair of variables (e.g. task condition and stimulus location), characterized by a pair of normalized weights, defines an angle (*ϕ*) in the plane spanned by the two variables. The distribution of these angles is compared with a null distribution obtained by shuffling the neuronal indexes. **f-h** Comparing the distribution of angles (green histograms) to the generated null distributions (orange), for time vs stimulus selectivity (**f**), time vs task selectivity (**g**), and task and stimulus selectivity (**h**). For each pair comparison, the half of the neurons with highest total joint-selectivity score were considered (Methods). All distributions are significantly different from the corresponding null distribution (Kolmogorov–Smirnov tests). The null distribution are not flat because the variable-informative dimensions were defined as non-linear combinations of dPCs (***Figure 3—figure Supplement 1***). The neurons falling into the extreme bins of the histogram in **h** are considered for the analysis in ***Figure 4***. **Figure 3—figure supplement 1**. Random distribution of selectivity between two dimensions.

To ensure the robustness of this result, we then applied the ePAIRS algorithm to selectivity spaces formed by a different number of dPCs ranging from 2 to 50 (***Figure 3c***). These dPCs cover up to 78.2 % of the total variance (***Figure 3d***). For spaces composed by 6 up to 23 dPCs, the outcome of the ePAIRS remains stable and the test indicates a significant degree of clustering in the full range from 2 up to 50 dPCs (all *p* values below 10^−9^). The decrease of the statistical measure of the test for space composed by more than 23 (***Figure 3c***) is likely due to the decreasing amount of task-relevant information of the subsequent dPCs and the corresponding increase in the amount noise. It is relevant that the population of PFC neurons shows significant structure for spaces down to a few number of dPCs, because it allowed us to gain some insight into the population structure based on the leading PCs that are directly linked to a task-relevant variable (***Figure 1***). Subsequent, less variance-explaining dPCs tend to capture mixtures of task-variables and their interpretation becomes more intricate (and in general high-dimensional structure is more difficult to analyze and interpret).

We thus focused on investigating the non-random, categorical encoding in the leading 6 dPCs, by considering each neuron’s contribution to combinations of task variables (***Figure 3e-h***). While this certainly cannot capture the full population structure present in the data, it enabled us to gain insight into the low-dimensional population structure for easily interpretable task-related features. We obtained a more detailed view of the non-random structure present in the data by considering each neuron’s contribution to a pair of features (***Figure 3e***; ***Yang et al., 2022***). The analysis is based on considering each neuron’s contribution to a pair of features as a two-dimensional vector and computing the angles that these vectors form with the x-axis in the respective two-dimensional feature space. The distribution of the angles in the data is then compared to null distributions obtained by bootstrapping (***Figure 3f-h***). We found that the original angle distribution is significantly different from the null distribution for all the two-feature comparisons: time-stimulus, time-task and task-stimulus (***Figure 3f-h***, Kolgomorov-Smirnov test all *p* < 0.005). Compared to the null distributions all the histograms show a slight concentration of neurons towards 0^°^ and 90^°^, indicating a slight over-representation of neurons with pure selectivity to one feature. Though the functional specialization to single features is higher than expected by chance, the histograms are not bimodal (according to a diptest), meaning that the neurons cannot be clearly classified into categorical groups. In other words, most of the neurons were characterized by intermediate angles and are thus best described as mixed selective.

### Task-selective functional groups respond differently during cue and delay epochs

Having established that most task-selective neurons are mixed selective to stimulus and task variables (***Figure 3h***), we wondered whether we could get some insight on the nature of this mixing. We found that these neurons form two categories with differential encoding of stimulus information during the delay period (***Figure 4***). To establish this result, we examined the weights of the first task-condition dPC (***Figure 4a***). These weights indicate how the neurons contribute with task condition-specific information or about how much a neuron’s activity is different between the memory and visual tasks. The distribution of the task-dPC weights shows heavier tails than a null distribution, obtained by shuffling the task labels (memory, visual) before the dPCA (***Figure 4a***). This suggests the presence of neurons with positive and negative non-zero weights, i.e., neurons that encode task-condition in *opposite* ways. It is important to note that the distribution in ***Figure 4a*** contains neurons with either stimulus selectivity, task selectivity or both. When we considered only neurons with mixed selectivity to task condition and stimulus, according to the analysis above (10^°^ < Φ < 80^°^ in ***Figure 3h***), the task-weight distribution was significantly bimodal (Figure 4b; diptest, *p* = 0.001; permutation test, *p* = 0.042).

**Figure 4.**
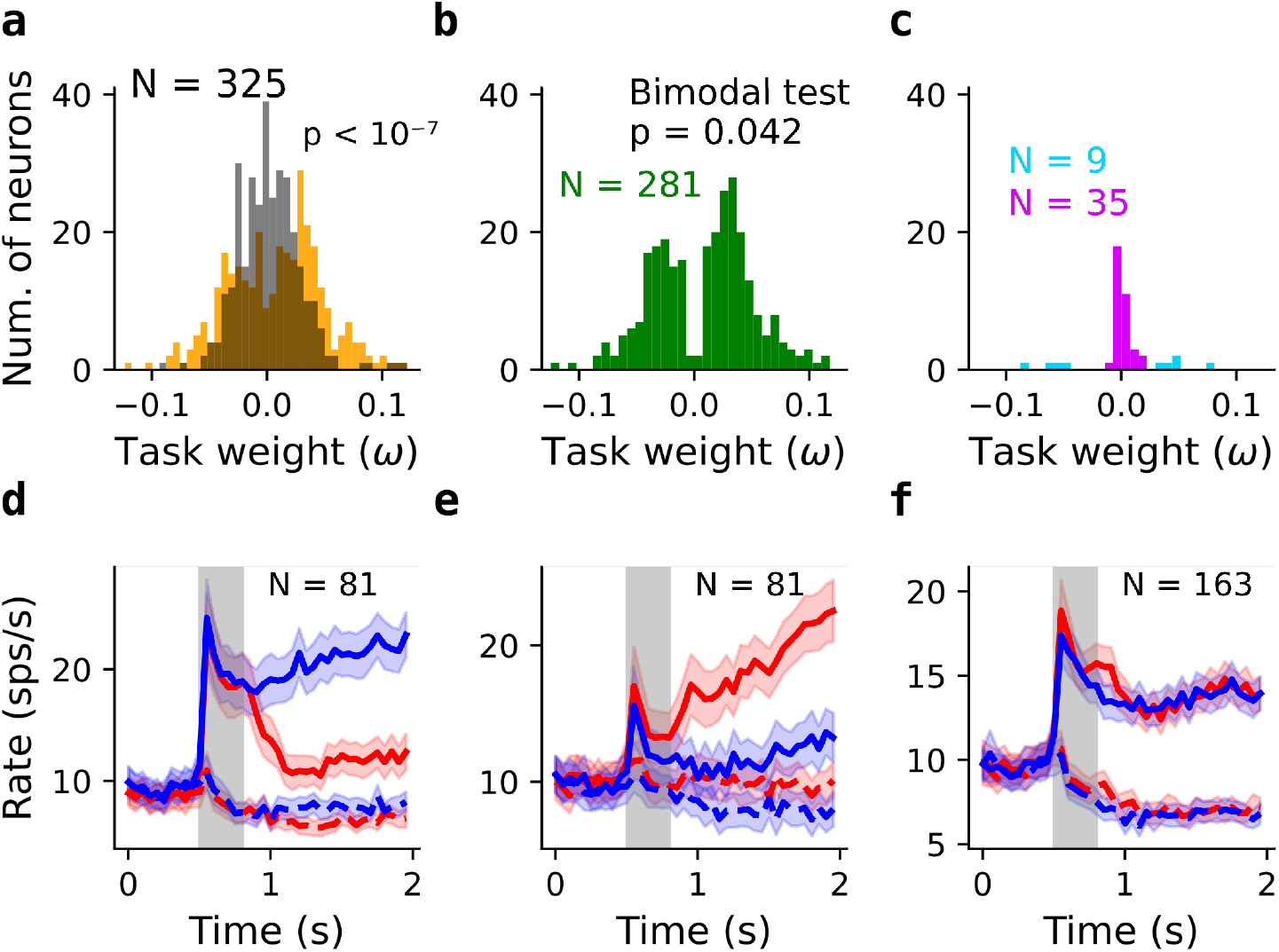
Functional subpopulations in mixed stimulus- and task-selective neurons. **a** Distribution of the task-dPC weights *ω* (orange) for the neurons with highest stimulus and task selectivity (***Figure 3e***) and null distribution when shuffling the task-condition labels before applying dPCA (gray). **b** Distribution of the task-dPC weights *ω* for neurons that encode a mixture of task and stimulus (green bins in ***Figure 3h***). The distribution is significantly bimodal (*p* = 0.042 after applying diptest to 10,000 random permutations of the original weights; ***Figure 4—figure Supplement 1***). **c** Distribtion of *ω* for pure stimulus-selective neurons (magenta bins in ***Figure 3h***) and pure task-selective neurons (cyan bins in ***Figure 3h***). **d-f** Average firing rates of three groups of neurons, defined according to their task-dPC weights *ω*. Red and blue differentiate memory from visual conditions, while solid and dashed indicate preferred and anti-preferred cues. **d** Neurons with task-dPC weight below the 25^*th*^ percentile (all *ω* < 0). **e** Neurons with task-dPC weight above the 75^*th*^ percentile (all *ω* > 0). **f** Remaining neurons with task-dPC close to zero. **Figure 4—figure supplement 1**. Bimodality test and firing rates of neurons with positive and negative task weights.

This segregation of mixed stimulus-task selective neurons in two distinct groups, with positive and negative weights, can be directly related to their function by visualizing their activity profiles ***Figure 3d-f***). Neurons with negative task weights show stimulus selective activity, but only when the stimulus is present on the screen (***Figure 4d*** and ***Figure 4—figure Supplement 1c***). These neurons increased their firing rates in response to their preferred cue location during the cue period, which is identical in the two tasks. In the delay period, their activity sharply decreased in the memory task but in the visual task it was maintained until stimulus offset at the end of the trial. Thus, this population of “encoding” neurons might play a role in stimulus encoding. In contrast, neurons with positive task weights showed stimulus selective activity predominantly in the delay period of the memory task, with increasing firing rates towards the end of the delay (***Figure 4e*** and ***Figure 4— figure Supplement 1d***). In the visual task, which does not require working memory, they did not activate. This activity profile of “mnemonic” neurons suggests that they might be implicated in memory maintenance. The apparent ramping activity may reflect time tracking (***Gouvêa et al., 2015***; ***Cueva et al., 2020***) or response anticipation (***Inagaki et al., 2019***).

Finally, as expected, the population of neurons with small absolute task weights did not show differential activation in the two task conditions and seemed to contribute to a stable representation of stimulus information during the memory delay independent of task context (***Figure 4d***). Taken together, this analysis revealed a categorical structure of the stimulus-task selectivity in the population of PFC neurons, suggesting a role of these populations in stimulus encoding and memory maintenance.

### Representation of cue and task information in neural population activity

In order to connect our findings regarding the geometry of low-dimensional population activity (***Figure 1h***) and the categorical mixed selectivity (***Figure 4***), we used the generalization performance of linear neural decoders across task conditions. Across task generalization is indicative of a shared low-dimensional representation across tasks (***Bernardi et al., 2020***). In particular, the similar two-dimensional geometry that we observed for the cue locations in the visual and memory tasks (***Figure 1h***) suggests that decoders trained on decoding the stimulus location in one task should generalize to the other task. We first confirmed that the cue location can be accurately decoded in each task when training and testing the decoder at trials from the same task using leave-one-out crossvalidation (***Figure 5a,b***; thick solid lines). For cross-task decoding (i.e., training on visual and testing on memory trials or vice versa), the accuracy during the cue period was as high as for the within-task decoders that were trained and tested in the same task (***Figure 5a,b***; dashed lines). However, during the delay period, across-task accuracy was weaker than within-task accuracy, although it stayed well above chance. This can be understood as a consequence of the functional specialization of neurons in encoding and mnemonic neurons (***Figure 4d,e***) that carry strong stimulus-related information during the delay period only in one of the two tasks. When training the decoder on separate conditions, it learned to rely more on the most informative neurons of the given task. The contribution of the neurons to the read-out is thus not optimal because the stimulus selectivity of neurons is task-dependent. However, the overall geometry of task-related information is preserved across tasks and this is why cross-task decoding is possible to some degree. To test whether it is possible to faithfully decode the stimulus location in a task-indepdent way, we trained a single decoder with trials from both tasks (with a matched number of trials to the previous decoding analysis). We found that this joint decoder performed as well as the within-task decoders (***Figure 5a,b*** thin lines). In other words, when trained with trials from both tasks, the decoder learned to readout stimulus location from an optimal task-independent projection of the neural activity onto a common stimulus-relevant plane. Taken together, the decoding results are consistent with both the low-dimensional task geometry for both tasks and the presence of separate functional populations with mixed selectivity for stimulus and task.

**Figure 5.**
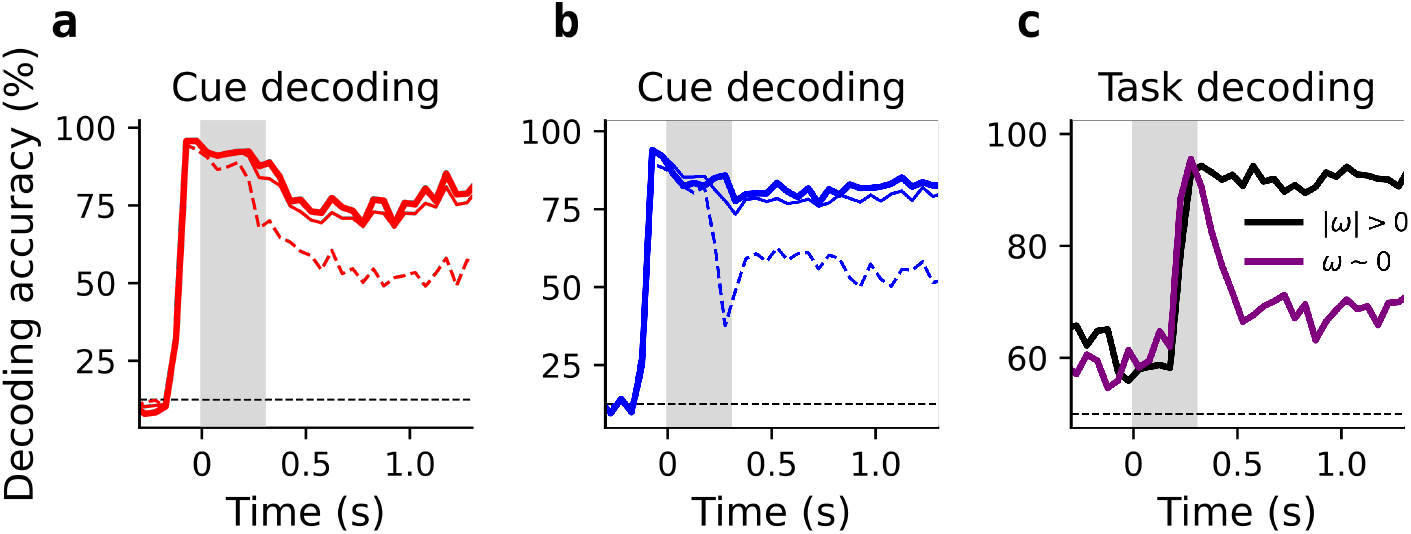
Decoding stimulus location and task condition reflects the structure of the neuronal representation. **a**,**b** Decoding the location of the presented cue using linear support vector machines with leave-one-out cross-validation (for the same N = 325 neurons as in ***Figure 4***). **a** Testing a decoder on memory trials, when trained on memory trials (thick solid line), memory and visual trials (thin solid line) and visual trials (dashed line). **b** The converse of **a**: testing a decoder on visual trials when trained on visual (thick solid line), visual and memory (thin solid line) and memory trials (dashed line). **c** Decoding task condition from the encoding and mnemonic neurons (***Figure 4d,e***, black line) and from the remaining, non-task selective, neurons.

Finally, we tested whether task condition can be reliably decoded from mnemonic and encoding neurons because they respond differently to the memory and visual context. Indeed, decoding accuracy was high in neurons with high task-condition dPC weights and only weak in the remaining neurons (***Figure 5c***). This result can also be related to the geometry of the representation: a decoder trained with encoding and mnemonic neurons learns to read out along the task-informative dimension (vertical task dPC axis in ***Figure 1h***). Contrarily, when using only the non-task-informative neurons, the representation of the neural trajectories are collapsed to a space that contains stimulus but not task information (horizontal stimulus plane in ***Figure 1h***), preventing reliable decoding of task condition. The finding that only a subset of the neurons carried task-related information provides further evidence for the presence of specialized functional subpopulations.

## Discussion

We have shown that in macaque prefrontal cortex stimulus and task information during working memory is encoded in a categorical, non-random way in neural subpopulations with different response profiles. Neuron with mixed selectivity for stimulus location and task condition show two contrasting activity profiles, with one population mostly responding during visual stimulation and the other population during memory maintenance. These subpopulations roughly correspond to the “early storage” and “late storage” neurons that were identified previously based on statistical tests of persistent activity on the single neuron level (***Markowitz et al., 2015***). Interestingly, these different functional cell types were concentrated at different anatomical locations, consistent with the idea that they may be part of different subnetworks (***Markowitz et al., 2015***). Our principled quantification of non-random structure in the distribution of feature selectivity among prefrontal neurons suggest that cortical populations responses in basic working memory and decision making tasks may share the same organization principles across primates and rodents (***Hirokawa et al., 2019***; ***Yang et al., 2022***).

We found that neural decoders generalize between memory and visual tasks, though with a decrease in performance compared to within-task decoders. Successful cross-task decoding is an indication of a low-dimensional neural representation (***Bernardi et al., 2020***), and it is consistent with the low-dimensional neural geometry suggested by the leading demixed principal components (***Figure 1h***). The lower performance of cross-task decoding compared to specialized within-task decoders is explained by the mixed selectivity of neurons to task and stimulus location which prevented the decoders to assign optimal weighting to neurons. Moreover, we found that a joint decoder, trained on data from both tasks, performed equally well as the specialized decoders. Thus, working memory information could be decoded in a task-invariant manner. Related to this is a study that found poor across-time generalization between time points before and after a distractor in a working memory task, unless the decoder was trained appropriately to identify a common subspace (***Parthasarathy et al., 2019***).

Recent modeling studies have investigated under what conditions trained recurrent neural networks (RNNs) learn representations relying on subpopulations or random mixed selectivity. ***Dubreuil et al. (2022***) showed that more complex tasks require a non-random population structure, in particular tasks that include flexible input–output mappings. Another important factor in RNN training is that the emergence of subpopulation structure depends on how the networks are initialized and also the optimization scheme (***Farrell et al., 2023***). Our tasks have a simple input-to-output mapping and could be solved by either networks with subpopulations or random structure. A further study (***Zhou et al., 2023***), found that complex and high-dimensional activity patterns, in the form of cue-selective sequences, emerged in tasks that require an explicit representation of time. Classical working memory tasks led to RNNs with low-dimensional population structure in which individual neurons show persistent and ramping activity, consistent with our results.

What can we infer from our results about the potential structure of working memory circuits in the prefrontal cortex? We found that neurons show mixed selectivity with distinct categorical subpopulations with similar firing patterns across neurons. Unlike in the case of random mixed selectivity, where neural representations are best studied at the level of population activity only (***Barak et al., 2013***; ***Mante et al., 2013***; ***Kaufman et al., 2014***; ***Stroud et al., 2023***; ***Tye et al., 2024***), in our case, different firing patterns are likely to contribute to the emergent properties of the network and individual activity profiles may be directly interpretable in terms of circuit mechanisms. A hypothetical network consistent with our data could be composed of two subcircuits, an encoding and a storage circuit. The encoding circuit would have relatively weak recurrent interactions, such that neurons are only active while a stimulus is presented, and activity fades upon stimulus removal. The storage circuit would have strong recurrent connectivity such that it shows attractor dynamics (***Compte et al., 2000***; ***Wimmer et al., 2014***). However, to account for the finding that memory neurons only become activity during working memory periods, some control mechanism must prevent the storage circuit from becoming active during stimulus periods. This could be achieved through feed-forward inhibition or a slowly ramping signal that would push the circuit into the attractor regime after cue offset. Functionally, such a circuit architecture could be beneficial for protecting working memory from distractors, as in in the case of binary working memory (***Murray et al., 2017***). It may also be useful in flexible routing of information from sensory to prefrontal circuits that play a key role in memory maintenance.

Our results add to a growing body of reports of disentangled, low-dimensional, representations of working memory information in primate prefrontal cortex (***Xie et al., 2022***; ***Lin et al., 2023***; ***Wu et al., 2020***). This opens the door to study the dynamics of neural circuits to gain mechanistic insights linking neural manifolds and underlying functional cell types.

## Methods and Materials

### Experimental paradigm

Full experimental details for the dataset can be found in ***Markowitz et al. (2015***). Two adult rhesus macaque monkeys (*Macaca mulatta*) were used for the study. The animals were trained for several weeks to perform a memory-guided saccade-task also called oculomotor-delayed-response or ODR task (***Fuster and Alexander, 1971***; ***Funahashi et al., 1989***; ***Constantinidis et al., 2001b***,a). In the ODR, which we refer to as memory task, the monkeys must first maintain their gaze on a fixating target during 0.5 *s*. Then, while the monkeys fixate, a cue stimulus is presented during 0.3 *s* on one out of eight locations on the screen. Removal of the stimulus gives way to the delay period, which for this experiment was varied between 1 and 1.5 *s*. The extinction of the fixation target indicated the animal to perform a saccade towards the remembered cue position.

In the visual variation of the ODR task (***Figure 1a***, bottom), which we refer to as visual task, the cue-stimulus is not removed from the screen until the switching off of the fixation target signals the animals to respond. Visual and memory trials were randomly interleaved without being cued to the monkey. Including the visual variation of the ODR task is an important feature of this dataset, which makes it possible to compare the neural activity between a condition requiring working memory and a condition with similar behavioral demands that does not require working memory.

### Neural recordings

Experimenters removed part of the animals’ skull to implant a 32-unit electrode (low-profile recording chamber Gray Matter Research, MT) into the arcuate cortex. The activity of isolated neurons was recorded while the monkeys performed up to 500 trials of randomly interleaved memory- and visually-guided delayed saccades to one of eight targets for a liquid reward. An infrared optical eye tracking system (120 Hz sampling monitored the animals’ eyes position. We included neurons in our analysis that had more than 5 correct trials for each cue location and task condition, as well as an average firing rate of at least 1 Hz during cue and delay periods. This left us with 650 neurons out of a total of 746. Note that previous analysis of the dataset was restricted to a subset of 365 positively tuned regular-spiking units (***Markowitz et al., 2015***).

### Targeted dimensionality reduction - dPCA

Similarly to Principal Component Analysis (PCA), demixed Principal Component Analysis (dPCA; ***Kobak et al., 2016***), finds linear combinations of the neurons’ activity that contain relevant information about the experimental task. The difference between both methods is that dPCA finds directions in neuronal space with maximal variance related to specific task features (or to combinations of task features). In this way, dPCA disentangles the contribution of different features to the total variance. Mathematically, the dPCs are obtained as the dimensions that minimize reconstruction error, as described by ***Horn and Johnson*** (***1985***) in the derivation of PC as a minimization problem. This minimization is performed with respect to an averaged or *marginalized* version of the data, which specifically contains variance related to the targeted feature or feature combination. The loss function is defined as:

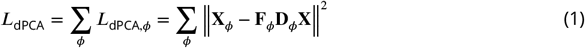

Where **X**_*ϕ*_ are the marginalizations of the data obtained as **X**_*ϕ*_ = **X** − _*ϕ*′≠*ϕ*_ *X* _*ϕ*′_ and **F**_*ϕ*_ and **D**_*ϕ*_ are the encoding and decoding matrices associated with feature *ϕ*. See ***Kobak et al***. (***2016***) for a detailed explanation and illustration of the method.

### Quantifying the degree of structured selectivity distribution with ePAIRS

Inspired by recent works (***Raposo et al., 2014***; ***Hirokawa et al., 2019***; ***Dubreuil et al., 2022***; ***Yang et al., 2022***), we used the “elliptical projection angle index of response similarity “(ePAIRS) to measure the presence of clusters in our data set. This test is a variation of the “projection angle index of response similarity “‘(PAIRS) method, which was first used in ***Raposo et al. (2014***). The method exploits the fact that neurons with similar selectivity will be represented as nearby points in a space spanned by feature-informative dimensions. A formal explanation follows.

Before applying the ePAIRS test, we reduced the dimensionality of our data. We show the results of applying the test to a different number of dPCs, but similar results are obtained using PCs. Applying dimensionality reduction is important to filter out the noise and irrelevant variance in the data and to obtain a set of feature dimensions we can interpret. We consider the matrix **D** whose *N* rows correspond to the number of considered neurons and *q* columns, the number of features (PCs or dPCs). Considering each row in **D** as a point in the *q*-dimensional feature space, the ePAIRS test compares the original distribution of directions **d**_*i*_/ ‖**d**_*i*_‖ to a null distribution which is obtained by bootstrapping from a multivariate Gaussian with the same covariance 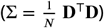 as the original data. The resampling ensures that the overall statistics of the data are maintained while population structure, if present, is disrupted. The algorithm is described by the following steps:

1. for each point **d**_*i*_ its *k* nearest neighbors are found (the points which maximize 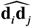). We have used *k* = 13 for our computations; smaller values of *k* do not alter the results qualitatively.
2. The mean *α*_*i*_ angle is computed for each neuron, defining the empirical distribution 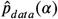.
3. Steps 1 and 2 are applied *N*_*bootstrap*_ times (we used *N*_*bootstrap*_ = 10, 000) to random distributions of *N q*-dimensional points, sampled from a multi-variate Gaussian *N*(0, Σ), to obtain the null distribution 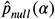
4. Finally, the original and null distributions are compared using a two-sided Wilcoxon rank-sum test, which yields a p-value. The effect size is computed as

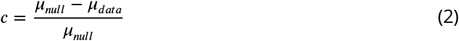
  - *c* > 0 means a smaller distance between neurons than expected by chance, i.e. structure or clusters.
  - *c* < 0 means greater distance than expected by chance, corresponding to neurons evenly spaced in feature space.

The difference between the ePAIRS and the PAIRS methods is that in ePAIRS, the null distribution is generated by bootstrapping from multivariate Gaussian distribution *with the same covariance* present in the data, while the PAIRS method samples from a spherical Gaussian distribution (the covariance matrix is diagonal with all elements in the diagonal having the same value). This difference can cause false positive results of the PAIRS method when applied to data with different variances along different features (which is usually the case, see ***Hirokawa et al***. (***2019***) for examples).

### Distribution of selectivity between pairs of variables

To get a finer-detailed view of how the neurons’ selectivity is distributed among different task features, we measured each neuron’s contribution to different pairs of activity modes. We used the same approach applied in ***Yang et al. (2022***) to assess the distribution of feature selectivity in the mouse ALM region. Each neuron’s contribution to a pair of features is represented as a two-dimensional vector in the space spanned by the corresponding features ***Figure 3—figure Supplement 1a***). The angle this vector forms with one of the axes measures how the neuron’s selectivity is distributed between the features. For every pair of features, the original distribution of angles is compared to a null distribution generated by bootstrapping over multivariate Gaussian distributions with data-matched variance. We conducted this analysis for combinations of time-stimulus, time-task, and task-stimulus selectivity, taking in each case half of the neurons from our dataset with the strongest combined selectivity (length of the corresponding feature vector) for the respective features. This selection is necessary because neurons that were not selective for either task variable would show a uniform random combination of features with very small vector lengths.

The feature-specific dimensions used in ***Yang et al. (2022***) were obtained directly as linear combinations in the space spanned by the neurons. They could define linear feature-specific dimensions because their task had two possible stimuli and two possible choices, so the corresponding stimulus and choice dimensions were one-dimensional. To analyze the contribution to pairs of features, the authors then used the absolute value of components or weights of each neuron along the corresponding feature vectors. For our analysis, we combined some of the feature-specific dPCs to obtain meaningful feature-specific dimensions. In particular, we combined the first two stimulus-dPCs because they provide complementary information related to variance along orthogonal directions on the plane where the stimuli were presented (see ***Figure 1***). In a similar way, we combined the first two time-dPCs, because they alone explained a substantial amount of the variance (14.5% and 7.3%). In both cases, the components of the combined dimensions are obtained as Euclidean norms:

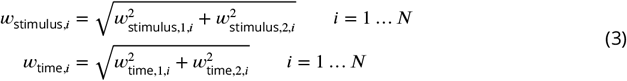

As a consequence of combining (non-linearly) dPCs to obtain the feature-specific vectors, the null distributions we obtain as controls are not flat. We show examples of null distributions of surrogate data in ***Figure 3—figure Supplement 1***.

### Classification according task dPC weight - quantiles

We classified the neurons into three functional groups based on their weights on the 1st task-dPC component. To obtain the three populations in ***Figure 4d-f*** we split according to the 25th and 75th percentile values.

### Bimodality tests

To determine whether the histograms of the weights in ***Figure 4b*** exhibited significant bimodality, we employed Hartigan’s dip test. This statistical measure calculates the maximum deviation between the observed distribution and the best-fitting unimodal distribution. It provides a statistical value *ν* and a corresponding significance *p*-value.

To ensure that the significance of bimodality was not solely due to the exclusion of certain neurons, we conducted a further analysis. We computed a *p*-value based on dip test statistics obtained from neurons selected using the same criterion (10^°^ < Φ < 80^°^), but after randomly shuffling the weights (10,000 random permutations). The resulting *p*-value was defined as the proportion of dip test statistics greater than that of the original data (***Figure 4—figure Supplement 1b***).

### Population decoding using support vector machines (SVM)

Classification algorithms that can decode the content of neural representations are often used to interpret data in cognitive and systems neuroscience (***Stokes et al., 2013***; ***Wolff et al., 2015***; ***Stokes, 2015***; ***King and Dehaene, 2014***; ***Kim et al., 2016***; ***Henderson et al., 2022***). They are particularly useful to identify the neural representations at the level of neural population activity. In general, a decoding analysis involves training an algorithm with a subset of the neural data and then testing its performance on the held out portion of the data.

Here, we used such a population decoding approach to quantify stimulus information at the population level. We trained and tested support vector machine (SVM) classifiers on surrogate populations of neurons (pseudopopulations) constructed from the individual neuron recordings (***Sarma et al., 2016***). In the pseudopopulations we included all neurons with more than 10 correct trials per condition (per task condition and stimulus combination). The spike count window was *T* = 50 ms. For each neuron, spike counts were normalized to zero mean and unit variance across all trials and time bins.

As classifiers, we used linear multi-class SVMs, C-SVC from LIBSVM (***Chang and Lin, 2011***). Non-linear SVMs with Radial Basis Function (RBF) kernels yielded the same results, so we used linear SVMs throughout the analysis. To estimate the decoder performance, we applied leave-one-out cross-validation. During each of 10,000 repeats, we selected a different set of 10 trials for each cue location for each neuron. Next, we randomly assigned 9 of the 10 trials to the training set and the remaining trials to the test set. This gave us a pseudopopulation response with 72 training trials and 8 test trials. For the joint decoder trained on data from both tasks, we used the same number of trials, randomly chosen from both tasks. The overall performance was calculated by averaging the test performance across all repetitions.

### Software and packages

To perform the dPC decomposition, we utilized the MATLAB functions developed by ***Kobak et al. (2016***). Similarly, for the ePAIRS test, we employed MATLAB functions adapted by ***Hirokawa et al. (2019***) from ***Raposo et al. (2014***). All subsequent analyses and figure generation were performed using Python.

## Acknowledgments

We acknowledge the use of Fenix Infrastructure resources, which are partially funded from the European Union’s Horizon 2020 research and innovation program through the ICEI project under the grant agreement No. 800858. This work was supported by grant PCI2020-112035 from MCIN/AEI/10.13039/501100011033 and the European Union ‘NextGenerationEU’/PRTR. This work was supported by the Spanish State Research Agency, through the Severo Ochoa and María de Maeztu Program for Centers and Units of Excellence in R&D (CEX2020-001084-M). We thank the CERCA Programme / Generalitat de Catalunya for institutional support.

---

**Figure 3—figure supplement 1.**
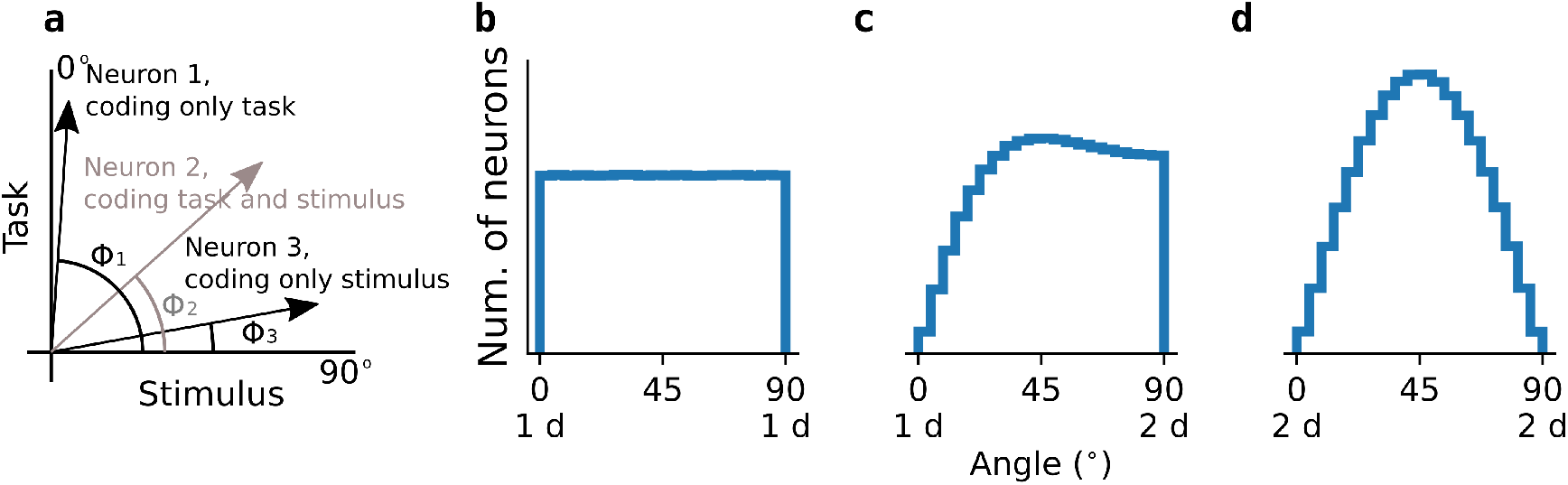
Random distribution of selectivity between two dimensions. **a** example selectivity of three cartoon neurons. **b-d** Null distribution of angles for a surrogate population of 1,000 neurons for different combinations of feature dimensions. **b** both dimensions are the absolute value of weights obtained as random samples from the same distribution. **c** one dimension is obtained as in **b**, the other dimension, as the square root of the sum of two squared weights, each obtained from samples of the same random distribution. **d** both dimensions are obtained as the second dimension in **c**.

**Figure 4—figure supplement 1.**
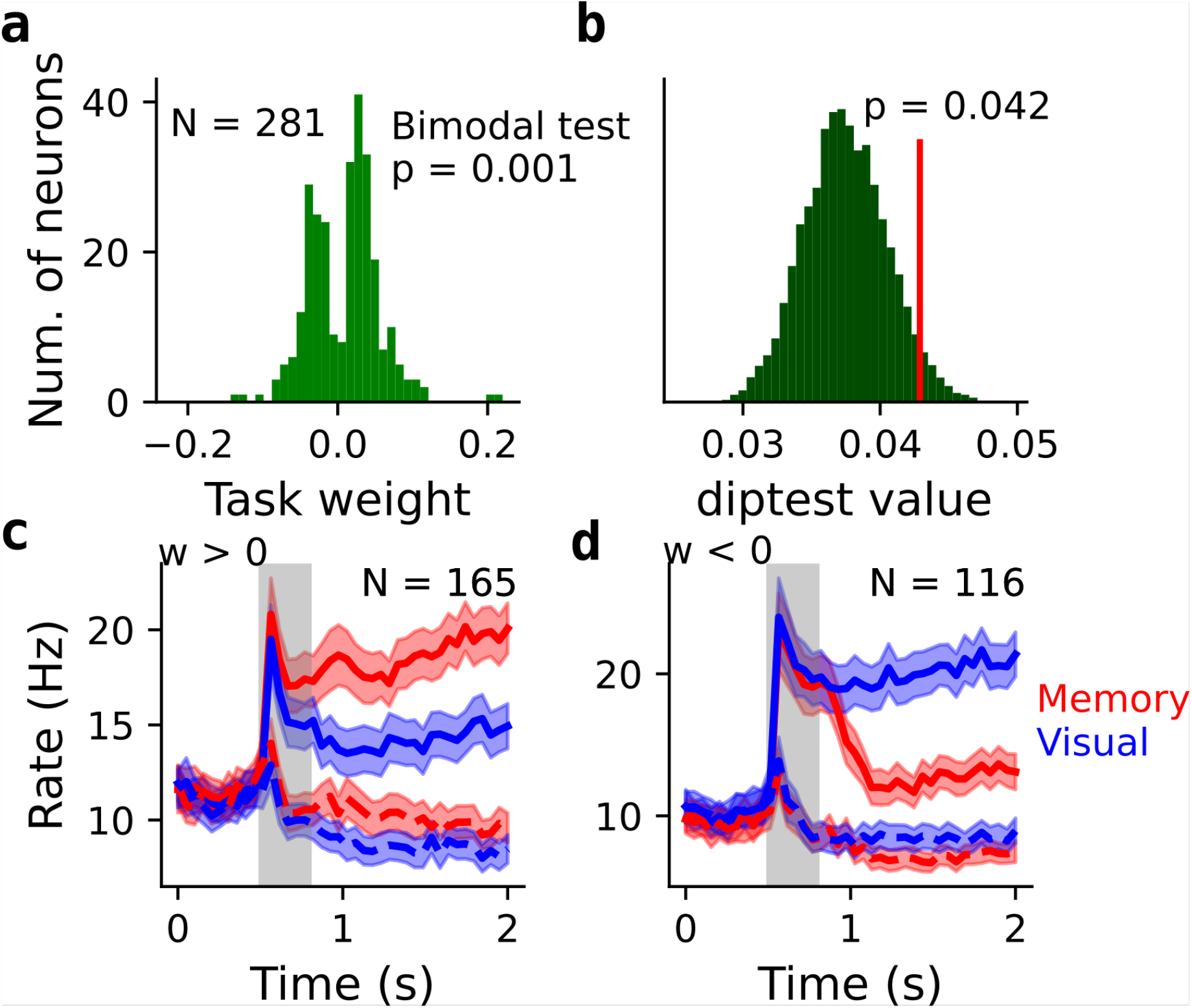
Bimodal contribution to task selectivity is related to contrasting activity profiles underlying encoding and maintenance processes. **a** Task dPC weight for the neurons with mixed task/stimulus selectivity (10^°^ < Φ < 80^°^ in ***Figure 3h***) showing a bimodal distribution (diptest *p* = 0.001) indicative of structured distribution of selectivity. **b** Test of significance obtained by comparing the diptest value of the original data to diptest value obtained when randomly shuffling the stimulus weights. The diptest values of 10,000 random shuffles were compared to the original diptest value, yieling *p* = 0.042. **c** Trial-averaged firing rate for the preferred (solid) and anti-preferred (dashed) stimulus locations during memory (red) and visual (blue) conditions for the PFC neurons with positive task weight. **e** same as **d** for the neurons with negative weight.

